## Supplementary material for "Detection of epistasis between *ACTN3* and *SNAP-25* with an insight towards gymnastic aptitude identification": The study protocol

**S1 Study Protocol:**

### Epistasis Detection in Genetic Data of Sports Gymnastics Contestants and Sedentary Individuals With an Insight Towards Classification Accuracy

This protocol has been provided by the authors to give readers additional information about the research work (.docx)

**Contents:**


1. **General Overview ____ 2**
2. **Confidentiality Issue and Data Censoring ____ 2**
3. **Confounders ____ 2**
4. **Inclusion / Exclusion Criteria ____ 2**
5. **STREGA Reporting Recommendations, Extended from STROBE Statement List ____ 3**

**References ____ 8**

#### 1. General Overview

This study has an observational case – control design. Gymnasts' representatives are extremely rare in any population. Especially, professional status is uncommon, e.g. in the Republic of Poland just 100 gymnasts compete at sub-elite sports try-outs (29 females, 71 males) and 96 contestants reached an elite level (44 females, 52 males) as described by Polish Gymnastics Federation in 2017.

#### 2. Confidentiality Issue and Data Censoring

The investigator ensured that a subject's confidentiality is maintained. On the case report forms or other documents mailed to the study office, subjects were identified by their initials and a subject study number only. Documents that were not for submission (e.g. signed informed consent forms) were kept in strict confidence by the investigator.

#### 3. Confounders

The influence of potential confounders were controlled via matching technique.

#### 4. Inclusion / Exclusion Criteria

The main inclusion criteria for individuals are as follows: male or female in the range of 16 - 28 years old, actual positive medical examination (regards sportsman), at least five years training experience and appropriate level of general and specific fitness (both points are eligibility criteria for sportsman), European ethnicity, elite or sub-elite sports level in gymnastics for cases, sedentary lifestyle for controls. In turn, exclusion points regard the data acquisition protocols. This means the maintenance of the standards required for DNA collection with storing and keeping high standards during surveying.

#### 5. STREGA Reporting Recommendations, Extended from STROBE Statement List

All the recommendations are considered in point-by-point overview either ad hoc or by referring to the main text of the manuscript.

1. (a) Title: Epistasis Detection in Sportsman Recognition Context: Gymnastics Matter. This study has an observational case – control design;

   (b) Information detailed in abstract;
2. Described in the introduction;
3. Described in the introduction; the study is the first report of genetic interaction study between performance enhancing polymorphisms (STREGA);

4. /5. Investigation procedure and critical paths

The study protocol was organized in two days. In the first one to obtain appropriate permission an investigator distributed official consents for participation in the study among adult athletes (> 18 years old) and/ or sportsman's parents or legal guards (participants < 18 years old). Afterward sportsman's health charts were supervised, if available. In parallel, the interview was performed in the form of an Athlete's Questionnaire or Control Group Representative Questionnaire. Especially, most important information regarded: age, nationality, years of training experience, amount of drills per week, frame of mind, social and economic conditions, drug use, dietary plan, relatedness issue, past injuries. Training drill was the only exposure factor that distinguished cases from controls.

The personal best achievements in official competitions were provided by coaches or the athletes themselves independently corroborated. Next, after seven days break the second part of the research procedure begun. This time DNA material was acquired with the help of saliva self-administered collection kits. Typically, saliva was taken early in the morning, before the first meal. Preliminary arrangement with coaches and managers let us specify <

100 elite and sub-elite male or female gymnasts of Republic of Poland, Lithuania and Italy, who were interested in supporting this project.

The research group members volunteered in Poland, Italy, Lithuania between 2011 and 2017. DNA isolation and genotyping were performed in the molecular laboratory of Gdansk University of Physical Education and Sport, Poland.

6. (a) Detailed in method section – participants description;

(b) The proportion of gymnasts to sedentary, healthy individuals was ca 1:4. Controls were matched to cases in the range of age: 16 – 28 yr. old, so that the difference between averages was less than 1 yr. old; sex proportion in cases and controls was assumed to differ less than 10 percentage points. All participants represented the same European ethnic group. Also social / economic conditions were the same in both groups and were reported as sufficient. The exposure factor regarded the sports regimen only.

(a) Variables are detailed in method section;

(b) All participants were unrelated European men (59.4%) or women (40.6%), and all of European descent (as self-reported) for ≥ 3 generations. Therefore, the influence of racial genetic skew has been minimized and the potential population stratification problems have been controlled.

SNPs being investigated were overviewed in introduction;

7. (a, b) The study is single-center research. Genotypes were assigned in batches (up to eighty-four 384-well plates). Regardless of the groups being genotyped, all sources of data and details of methods of assessment (measurement) is given method chapter;

8. (a, b) The bias was controlled by eligibility criteria;

9. Participants and sample size

The study was performed on sportsmen and sedentary individuals. Information on the prevalence / occurrence of gymnasts in the population was obtained from the Polish

Gymnastics Federation and Central Statistical Office [1-2]. In 2016, 3871 gymnasts were registered and 196 of them comprised elite and sub-elite athletes. Furthermore, the range of legitimized sportsman across voivodeships in Poland vary between 18 and 36 instances. Thereafter, the formula to estimate the proportion of gymnast to non-gymnast in a dichotomous outcome variable in a single population is as follows [3]:

$$N = probability(1 - probability)\left(\frac{1{,}96}{0{,}05}\right)^2.$$

Assuming: p-value= 0.05, 36 sportsmen per 1000 inhabitants, and 10% dropout of athletes from the research program, the sample size of N= 55 satisfy the above formal calculation. Because of lower than expected event rate, it was of significant demand to enroll up to 20% gymnasts to the whole research group;

10. Quantitative variables were analyzed in categories. Naturally, data were processed for sportsmen and controls separately in descriptive statistics. For multidimensional computations, binary outcome were set;

12. (a and b, d – i) Information specified in method section;

(c) Handling of Missing Data

In the analysis, missing data were not considered. Though, imputation techniques might bring several benefits, e.g. increased precision of the estimates and potential biases reduction.

(j) Relatedness

Samples were drawn from large, non-isolated populations. None of the individuals had any connections and were not distant relatives. However, the total number of participants pairs was (N*N-1)/2. After 318 randomized draws of i, j instances pairs, the Similarity Index of [4] formula was adapted to calculate z-score statistics form the general pattern ($\frac{observed - expected}{SD\ observed}$). Next bootstrap technique was applied to obtain 52000 sets of pairs. According to the data that was generated, we concluded participants enrolled in the study are unrelated at p-Value < .0001;

(g) Inferring genotypes

To gather the molecular data PCR and RT-PCR techniques have been applied. Genotypes were inferred following ethidium bromide staining of 2% agarose gel and exposure to UV light.

*ACTN3* (*rs1815739*) genotyping

The 290 bp fragment of exon 15 of the *ACTN3* gene was amplified using the forward primer: 5’- CTGTTGCCTGTGGTAAGTGGG-3’ and the reverse primer: 5’-TGGTCACAGTATGCAGGAGGG-3’. The alleles *577R* and *577X* were distinguished by the presence (*577X*) or absence (*577R*) of a *Dde I* restriction site (Thermo Fisher Scientific, Poland). Digestion of PCR products of the *577X* allele yields bands of 108 bp, 97 bp and 86 bp, whereas digestion of PCR products of the *577R* allele yields band of 205 bp and 86 bp.

*PPARGC1A* (*rs8192678*) genotyping

The *PPARGC1A* polymorphism was amplified as follows: forward 5’-TTGTTCTTCCACAGATTCAGAC-3’ and 5’ -GAAAAGGACCTTGAACGAGAG-3’ (reverse primer). The analyzed PCR product was cut by *Msp I* (Thermo Fisher Scientific, Poland) restriction endonuclease (Thermo Fisher Scientific, Poland). Digested PCR fragments were of 449 bp length for the *G* allele and 274 bp or 175 bp for the *A* allele.

*PPARα* (*rs4253778*) genotyping

The intron 7 polymorphism (*rs4253778*) was studied, using a 5′ forward primer of ACAATCACTCCTTAAATATGGTGG and AAGTAGGGACAGACAGGACCAGTA (a 3′ reverse primer). The PCR products were incubated (digested) with the restriction enzyme *Taq1* (Thermo Fisher Scientific, Poland) which cleaved the amplicon (266 bp) from the *C* allele into two fragments of 216 bp and 50 bp length.

*BDNF-AS* (*rs6265*) genotyping

The primers were designed to amplify a fragment of length 268 bp, which contains the site of the SNP. Forward: 5’-TGACCCACTTGCCACCCGTGC-3’ and reverse: 5’-GCAGCAGCCAGGGCTGGC-3’ primers were used for amplification followed by cutting with *BsaJ1* restriction enzyme (Thermo Fisher Scientific, Poland). Alleles *T* represent the absence of restriction site (268-bp), while alleles *C* indicate the presence of restriction site (152 bp and 116 bp bands).

*GNB3* (*rs5443*) genotyping

The forward primer was 5′-TGACCCACTTGCCACCCGTGC-3′, and the reverse primer was 5′-GCAGCAGCCAGGGCTGGC-3′. A 25 base pairs DNA marker was used, it ranged from 25 to 300 bp. The *CC* genotype gave two bands of 115 and 152 bp, *CT* was expected to give three bands of 267, 152, and 115 bp, while *TT* gave one band of 267 bp. The PCR products were incubated with the restriction enzyme *BsaJ1* (Thermo Fisher Scientific, Poland).

*DRD2* (*rs1076560*) genotyping

Amplification of the 213 bp DNA fragment containing the *rs1076560* polymorphism (*G>T*) was performed using forward: 5′-GGCAGAACAGAAGTGGGGTA-3′ and reverse: 5′-GACAAGTTCCCAGGCATCAG-3′ primers. Amplified PCR products were digested with 1 unit of *Hph1* enzyme (Thermo Scientific, Warsaw, Poland).

*SNAP-25* (*rs362584*) genotyping

Genotyping of *SNAP-25 – rs362584* located in the flanking region was conducted using TaqMan assays C_330323_20 (Thermo Scientific, Warsaw, Poland; Catalog number: 4351379), including primers and fluorescently labeled (Context Sequence [VIC/FAM]: ACAAGAGCCCATGCCTTCTTCAAAT**[A/G]**TGCTACTCACAGAGAATTTTTCATT) MGB probes for detection of the alleles.

13. Genotyping was successful in 318 individuals;

14. – 16. (a, d), 17. (a, c) Information is presented in the Results section;

18. – 22. These points are summarized in Discussion and Other Information.
