## Supplemental text and tables for "Detection of epistasis between *ACTN3* and *SNAP-25* with an insight towards gymnastic aptitude identification"

**S2 Supplementary Material:**

### Epistasis Detection in Genetic Data of Sports Gymnastics Contestants and Sedentary Individuals With an Insight Towards Classification Accuracy

This work contains all supplemental text, figures, and tables. (.docx)

**Contents**

1. **Summary of The Results From Table S1 ____ 2**
2. **Table S1 ____ 2**
3. **Table S2 ____ 6**
4. **Table S3 ____ 7**
5. **Table S4 ____ 7**
6. **Table S5 ____ 7**
7. **Table S6 ____ 7**
8. **Coding schemes applied to data analysis ____ 8**
9. **Genotypes numbering applied to simple linear coding with literature support ____ 8**

**References ____ 9**

#### 1. Summary of The Results From Table S1

A compiled calculations for statistically significant intercepts and cross-partial G-G interactions detected with additive – multiplicative logistic regression models through eighteen coding data arrangement on one reference category and an example, has been given in Table S1. Interpretation of the coefficients is not straightforward, and in deep meaning depends on null hypothesis as well as on data arrangement. As an example of *ACTN3* – *SNAP*-25 and orthogonal data coding, $b_{RR,GA\,(1,2)}$ = 1.564 (p-value = 0.1) weight affects positively on 2$^{nd}$ – 2$^{nd}$ carriers and on 2$^{nd}$ – ref. genotype it influences in reveres way. Consequently, $b_{RR,GA}$ itself promotes ref. – ref. gymnasts much more than 2$^{nd}$ – 2$^{nd}$ and 2$^{nd}$ – heterozygous representatives: 22.8 (95.8%) to 4.8 (82.7%) in the sense of odds ratio and probability for both comparisons. Moreover, $b_{(2,2)}$ interaction term (orthogonal contrasts), despite being positive, reduces odds of success in gymnastics in *XX* – *AA* contestants to 0.044 (4.2%). Interestingly, $b_{(2,2)}$ interaction weight of *ACTN3* – *SNAP-25* is always negative and significant between 0.03 to 0.09, regardless the type of coding except for reverse Helmert data arrangement (P = 0.12), and should be understand as highly informative in gymnastics. As important seems to be $b_0$ weight in G – G interaction for *ACTN3* – *SNAP-25*, *PPARα* – *PPARGC1A* and *GNB3* – *DRD2*, when reference-specific coding is assumed (Table S1). Here, in very particular context, the genotypes may disfavour athletes in strength / power specialities. Ultimately, standard linear scheme vs. genetic linear approach gives spurious results due to *PPARGC1A* – *SNAP-25*, *BDNF* – *GNB3*. For this reason, in-depth review of cross-partial dependencies is indispensable. Nonetheless, linear coding in both: 1, 2, 3 coding (simple one) and genetic coding are consistent for *ACTN3* – *PPARα* effect**.**

**2. Table S1. The summary for statistically significant intercepts and cross-partial G – G interactions detected with logistic regression through 18 coding schemes.**

| | (1) Dummy variable arrangement: |
|---|---|
| *ACTN3* – *SNAP-25*<br>(zero category: *XX*, *AA*) | $b_0$ = -2.71; P = 0.1,<br>$b_{RR,GG\,(2,2)}$ = -3.890; P = 0.09 |

| | |
|---|---|
| (2) Effect coding: | |
| *PPARα – PPARGC1A* (zero category: GG / *glygly*) | $b_0$ = -1.386; P = 0.004, $b_{CC,serser\ (2,2)}$ = -3.068; P = 0.002 |
| *ACTN3 – SNAP-25* (negative category: *XX*, *AA*) | $b_0$ = -1.152; P = 0.02, $b_{RR,GG\ (1,1)}$ = -1.233; P = 0.07, $b_{RX,GA\ (2,2)}$ = -1.229; P = 0.05 |
| *PPARGC1A – SNAP-25* (negative category: glygly, *AA*) | $b_0$ = -1.188; P = 0.02, $b_{glyser,RR\ (2,1)}$ = 1.224; P = 0.06, $b_{glyser,GA\ (2,2)}$ = -0.975; P = 0.08 |
| *PPARGC1A – GNB3* (negative category: glygly, *TT*) | $b_0$ = -1.110; P = 0.02, $b_{glyser,CC\ (2,1)}$ = 1.129; P = 0.06, $b_{glyser,TC\ (2,2)}$ = -0.593; P = 0.02 |
| (3) Orthogonal 1 (2nd), 0, -1 (ref.) vs. 1 (2nd), 1, -2 – (ref.) coding: | |
| *ACTN3 – SNAP-25* (reference category: *XX*, *AA*) | $b_0$ = -1.152; P = 0.02, $b_{RR,GG\ (1,1)}$ = -3.591; P = 0.03, $b_{RR,GA\ (1,2)}$ = 1.564; P = 0.1, $b_{RX,GG\ (2,1)}$ = 2.023; P = 0.04, $b_{RX,GA\ (2,2)}$ = -1.229; P = 0.05 |
| *PPARGC1A – SNAP-25* (reference category: glygly, *AA*) | $b_0$ = -1.189; P = 0.02, $b_{glyser,GG\ (2,1)}$ = 2.199; P = 0.04, $b_{glyser,GA\ (2,2)}$ = -0.975; P = 0.08 |
| *PPARGC1A – GNB3* (reference category: glygly, *TT*) | $b_0$ = -1.110; P = 0.02, $b_{glyser,CC\ (2,1)}$ = 1.723; P = 0.05, $b_{glyser,TC\ (2,2)}$ = -0.593; P = 0.01 |
| (4) Dummy variables polynomial – orthogonal coding for linear (1, 0, -1 – reference) vs. quadratic (-1, 2 – heterozygous, -1 – ref.) trend: | |
| *ACTN3 – SNAP-25* (reference category: *XX*, *AA*) | $b_0$ = -1.236; P = 0.02, $b_{(quadratic),(quadratic)\ (2,2)}$ = -0.318; P = 0.05 |
| *PPARGC1A – SNAP-25* (reference category: glygly, *AA*) | $b_0$ = -1.189; P = 0.02, $b_{(linear),(quadratic)\ (1,2)}$ = -0.499; P = 0.1, $b_{(quadratic),(quadratic)\ (2,2)}$ = -0.244; P = 0.08 |
| (5) Orthonormal-orthogonal coding with values of first variable: 0.5, 0.5 – heterozygous, ≈ -0.71 – reference and second variable: ≈ 0.71, ≈ -0.71 – heterozygous, 0 – reference: | |
| *ACTN3 – SNAP-25* (reference category: *XX*, *AA*) | $b_0$ = -1.199; P = 0.03, $b_{RX,GA\ (2,2)}$ = -1.795; P = 0.03 |
| *PPARGC1A – SNAP-25* (reference category: glygly, *AA*) | $b_0$ = -1.189; P = 0.03, $b_{serser,GA\ (1,2)}$ = 3.117; P = 0.04 |
| (6) Simple contrast coding and values of first variable: 0.66, -0.33 – heterozygous, -0.33 – reference, and second variable: -0.33, 0.66 – heterozygous, -0.33 – reference: | |
| *ACTN3 – SNAP-25* (reference category: *XX*, *AA*) | $b_0$ = -1.152; P = 0.02, $b_{RX,GA\ (2,2)}$ = -3.889; P = 0.09 |

| | |
|---|---|
| (7) Helmert contrast coding and values of first variable: 0.66 – dominant, -0.33, -0.33 – recessive, and second variable: 0 – dominant, 0.5, -0.5 – recessive: | |
| *ACTN3 – SNAP-25* (reference category: *XX*, *AA*) | $b_0$ = -1.152; P = 0.02, $b_{RR,GG\ (1,1)}$ = -2.774; P =0.07, $b_{RX,GA\ (2,2)}$ = -3.890; P = 0.09 |
| *PPARGC1A – SNAP-25* (reference category: glygly, *AA*) | $b_0$ = -1.189; P = 0.02, $b_{glyser,GG\ (2,1)}$ = 4.927; P = 0.05 |
| (8) Reverse Helmert contrast coding through values of first variable: -0.5, 0.5 – heterozygous, 0 – (ref.), and second variable: -0.33, -0.33 - heterozygous, 0.66 – (ref.): | |
| *ACTN3 – SNAP-25* (reference category: *XX*, *AA*) | $b_0$ = -1.152; P = 0.02, $b_{RR,GG\ (1,1)}$ = -3.591; P = 0.03 |
| (9) Mathematical genetic coding through values of first variable: -1, 0 – heterozygous, -1 – reference, and second variable: 0, 1 – heterozygous, 0 – reference: | |
| *ACTN3 – SNAP-25* (reference category: *XX*, *AA*) | $b_0$ = -1.604; P = 0.06, $b_{RX,GA\ (2,2)}$ = -2.902; P = 0.05 |
| (10) Self-defined coding through values of first variable: 1, 0 – heterozygous, -1 – reference, and second variable: 1, -0.5 – heterozygous, -0.5 – reference: | |
| *ACTN3 – SNAP-25* (reference category: *XX*, *AA*) | $b_0$ = -1.151; P = 0.02, $b_{RR,GG\ (1,1)}$ = -3.889; P =0.08, $b_{RR,GA\ (1,2)}$ = 4.242; P = 0.06, $b_{RX,GA\ (2,2)}$ = -4.912; P = 0.05 |
| (11) Forward difference coding through values of first variable: 0.66, -0.33 – heterozygous, -0.33 – (ref.), and second variable: 0.33, 0.33 – heterozygous, -0.66 – (ref.): | |
| *ACTN3 – SNAP-25* (reference category: *XX*, *AA*) | $b_0$ = -1.152; P = 0.02, $b_{RR,GG\ (1,1)}$ = -3.591; P = 0.04, $b_{RX,GA\ (2,2)}$ = -3.889; P = 0.09 |
| (12) Backward difference coding through values of first variable: -0.66, 0.33 – heterozygous, 0.33 – (ref.), and second variable: -0.33, -0.33 – heterozygous, 0.66 – (ref.): | |
| *ACTN3 – SNAP-25* (reference category: *XX*, *AA*) | $b_0$ = -1.152; P = 0.02, $b_{RR,GG\ (1,1)}$ = -3.591; P = 0.04, $b_{RX,GA\ (2,2)}$ = -3.890; P = 0.09 |
| (13) Repeated contrast coding through values of first variable: 0, -1 – heterozygous, -1 – reference, and second variable: 0, 0 – heterozygous, -1 – reference: | |
| *ACTN3 – SNAP-25* (reference category: *XX*, *AA*) | $b_0$ = -2.175; P = 0.03, $b_{RR,GG\ (1,1)}$ = -3.591; P = 0.03, $b_{RX,GA\ (2,2)}$ = -3.890; P = 0.09 |
| (14) Non-orthogonal coding through values of first variable: -2, 1 – heterozygous, 1 – reference, and second variable: -1, 0 – heterozygous, 1 – reference: | |
| *ACTN3 – SNAP-25* (reference category: *XX*, *AA*) | $b_0$ = -1.152; P = 0.02, $b_{RR,GG\ (1,1)}$ = -1.229; P = 0.05, $b_{RX,GA\ (2,2)}$ = -3.890; P = 0.09 |

| | |
|---|---|
| (15) Deviation coding and recessive – recessive treated as reference category with values of first variable: 0.5, 0 – heterozygous, -0.5 – reference, and second variable: 0, 0.5 – heterozygous, -0.5 – reference: | |
| *ACTN3 – SNAP-25* (reference category: *XX*, *AA*) | $b_0$ = -1.150; P = 0.02, $b_{RR,GG\ (1,1)}$ = -4.953; P = 0.05, $b_{RX,GA\ (2,2)}$ = -4.883; P = 0.07 |
| *PPARGC1A – SNAP-25* (reference category: glygly, *AA*) | $b_0$ = -1.183; P = 0.02, $b_{glyser,GG\ (2,1)}$ = 4.895; P = 0.06, $b_{2glyser,GA\ (2,2)}$ = -3.902; P = 0.08 |
| *PPARGC1A – GNB3* (reference category: glygly, *TT*) | $b_0$ = -1.110; P = 0.02, $b_{glyser,CC\ (2,1)}$ = 4.517; P = 0.06 |
| (16) Simple / standard linear coding for the 1, 2, 3 values[1]: | |
| *ACTN3 – PPARα* | $b_{ACTN3,PPAR\alpha}$ = 1.874; P = 0.05 |
| *PPARGC1A – SNAP-25* | $b_{PPARGC1A,SNAP\text{-}25}$ = 1.043; P = 0.07 |
| *PPARα – SNAP-25* | $b_{PPAR\alpha,SNAP\text{-}25}$ = 1.300; P = 0.08 |
| *PPARGC1A – SNAP-25* | $b_{PPARGC1A,SNAP\text{-}25}$ = 1.043; P = 0.07 |
| *BDNF – GNB3* | $b_{BDNF,GNB3}$ = -1.708; P = 0.06 |
| *GNB3 – SNAP-25* | $b_{GNB3,SNAP\text{-}25}$ = -1.172; P = 0.06 |
| *DRD2 – SNAP-25* | $b_{DRD2,SNAP\text{-}25}$ = -1.583; P = 0.05 |
| (17) Linear genetic coding[2]: | |
| *ACTN3 – PPARα* | $b_{ACTN3,PPAR\alpha}$ = 2.412; P = 0.04 |
| *PPARGC1A – SNAP-25* | $\mathrm{b_{PPARGC1A,SNAP\text{-}25}}$ = -3.461; P = 0.06 |
| *BDNF – GNB3* | $b_{BDNF,GNB3}$ = 2.548; P = 0.05 |
| (18) GLM coding with the same beta weight as in reference dummy coding: | |
| | The main difference concerns replacement of joint test to type 3 test joint test. Consequently, main effects were non-significant at P ≤ 0.05 in any reasonably G – G interaction. |

One out of three possible reference category adjustment for specific genotype coding is presented.

[1] The detailed description is given in chapter 9 on page 8.

[2] Genotypes were arranged according to information form Table 1 and Table S2.

**3. Table S2. The quality measures for seven investigated genetic markers.**

| Group / SNP's quality control | Gymnasts $n_1$= 73 | Controls $n_2$= 245 |
|---|---|---|
| *ACTN3* – rs1815739 | | |
| *XX* (%): | 25 (34.2) | 41 (16.7) |
| *RX* (%): | 29 (39.7) | 118 (48.2) |
| ancestral *RR* (%):(ref. R allele: R>X / C>T) | 19 (26.1) | 86 (35.1) |
| MAF *X* allele: | 54.1% | 40.8% |
| HWE / P-Value: | 2.922 / 0.09 | 0.002 / 0.961 |
| *PPARGC1A* – rs8192678 | | |
| *TT* (%): | 14 (19.2) | 62 (25.3) |
| *TC* (%): | 31 (42.5 | 103 (42.0) |
| *CC* (%): (ref. C allele: C>T / G>A) | 28 (38.3) | 80 (32.7) |
| MAF *T* allele: | 40.4% | 46.3% |
| HWE / P-Value: | 1.021 / 0.312 | 5.857 /0.016* |
| *PPARα* – rs4253788 | | |
| *CC* (%): | 9 (12.3) | 5 (2.0) |
| *CG* (%): | 28 (38.4) | 71 (29.0) |
| *GG* (%): (ref. G allele: G>C / G>T) | 36 (49.3) | 169 (69.0) |
| MAF *C* allele: | 31.5% | 16.5% |
| HWE / P-Value: | 0.904 / 0.342 | 0.616 / 0.433 |
| *BDNF-AS* – rs6265 | | |
| *TT* (%): | 6 (8.2) | 9 (3.7) |
| *TC* (%): | 24 (32.9) | 79 (32.2) |
| *CC* (%): (ref. C allele: C>T / G>A) | 43 (58.9) | 157 (64.1) |
| MAF *T* allele: | 24.7% | 19.8% |
| HWE / P-Value: | 0.968 / 0.325 | 0.058 / 0.809 |
| *GNB3* – rs5443 | | |
| *TT* (%): | 31 (42.5) | 24 (9.8) |
| *TC* (%): | 26 (35.6) | 106 (43.3) |
| *CC* (%): (ref. C allele: C>T) | 16 (21.9) | 115 (46.9) |
| MAF *T* allele: | 60.3% | 31.4% |
| HWE / P-Value: | 4.794 / 0.029* | 0.004 / 0.953 |
| *DRD2* – rs1076560 | | |
| *TT* (%): | 5 (6.9) | 8 (3.3) |
| *TG* (%): | 29 (39.7) | 73 (29.8) |
| *GG* (%):(ref. C allele: C>A / G>T) | 39 (53.4) | 164 (66.9) |
| MAF *T* allele: | 26.7% | 18.2% |
| HWE / P-Value: | 0.016 / 0.901 | 0.001 / 0.972 |

| | | |
|---|---|---|
| *SNAP-25 – rs362584* | | |
| *AA* (%): | 9 (12.3) | 29 (11.8) |
| *GA* (%): | 27 (37.0) | 103 (42.0) |
| ancestral *GG* (%):(ref. G allele: G>A) | 37 (50.7) | 113 (46.2) |
| MAF *A* allele: | 30.8% | 32.9% |
| HWE / P-Value: | 1.285 / 0.257 | 0.545 / 0.460 |

MAF is referenced due to the data from 1000 Genome Project.
* Significant at $p \leq 0.05$.

4. **Table S3. IF – THEN rules.**

| IF – THEN PARADIGM | CLASSIFICATION RULE |
|---|---|
| if *ACTN3* = 3 and *SNAP-25* = 2 | **1** |
| if *ACTN3* = 3 and *SNAP-25* = 3 | **0** |
| if *ACTN3* = 3 and *SNAP-25* = 1 | **1** |
| if *ACTN3* = 2 and *SNAP-25* = 2 | **0** |
| if *ACTN3* = 2 and *SNAP-25* = 3 | **1** |
| if *ACTN3* = 2 and *SNAP-25* = 1 | **1** |
| if *ACTN3* = 1 and *SNAP-25* = 2 | **1** |
| if *ACTN3* = 1 and *SNAP-25* = 3 | **0** |
| if *ACTN3* = 1 and *SNAP-25* = 1 | **0** |

5. **Table S4. Contingency matrix of genotype counts for *ACTN3 – rs1815739* and *SNAP-25 rs362584*.**

| *X1* | *X2* | | | Total |
|---|---|---|---|---|
| | 1 | 2 | 3 | |
| 1 | 16 | 29 | 10 | 55 |
| 2 | 12 | 70 | 67 | 149 |
| 3 | 5 | 24 | 49 | 78 |
| Total | 33 | 123 | 126 | 282 |

6. **Table S5. Basic statistics obtained on training data for *ACTN3 – PPARGC1A – PPARα – SNAP-25* epistatic model.**

| BAL. ACC. | ACC. | SENSIT. | SPECIF. | OR / CI | $\chi^2$ | $\chi^2$ p-val. | PRECIS. | KAPPA | F |
|---|---|---|---|---|---|---|---|---|---|
| 0.788 | 0.733 | 0.889 | 0.687 | 17.522/ 7.302 ; 42.048 | 59.724 | >0.0001 | 0.454 | 0.430 | 0.601 |

7. **Table S6. Basic statistics from whole dataset for *ACTN3 – PPARGC1A – PPARα – SNAP-25* epistatic model.**

| BAL. ACC. | ACC. | SENSIT. | SPECIF. | OR / CI | $\chi^2$ | $\chi^2$ p-val. | PRECIS. | KAPPA | F |
|---|---|---|---|---|---|---|---|---|---|
| 0.785 | 0.727 | 0.891 | 0.679 | 17.216/.471 ; 39.676 | 64.830 | < 0.0001 | 0.449 | 0.423 | 0.597 |

#### 8. Coding schemes applied to data analysis

Apart from fundamental, molecular types of genotypes ordering, e.g. additive, recessive, dominant, over-dominant, multiplicative, we recognized nineteen classic (statistical and mathematical) notations to describe SNPs. These are: (1) dummy, (2) effect, (3) orthogonal, (4) weighted effect, (5) polynomial, (6) orthonormal, (7) simple contrast, (8) Helmert contrast, (9) reverse Helmert contrast, (10) mathematical-genetic, (11) self-defined, (12) forward difference, (13) backward difference, (14) repeated contrasts, (15) deviation contrasts, (16) non-orthogonal coding, (17) a simple linear coding, (18) linear genetic coding, (19) GLM coding. Moreover, when three categories are distinguished, a three parallel, so called: reference categories, for each genotype (except for GLM design) can be applied.

#### 9. Genotypes numbering applied to simple linear coding with literature support

*ACTN3* – *rs1815739*

1: *XX* / 3: *RR* (ancestral) – 1: *TT* / 3: *CC* (ancestral)

*ACTN3 gene is involved in muscle contractions* [1].

*PPARGC1A* – *rs8192678*

1: *TT(Gly)* / 3: *CC(Ser)* – 1: *GG(Gly)* / 3: *AA(Ser)*

*Other studies, however, report the Ser allele as useful in power activities* [2].

*PPARα* – *rs4253788*

1: *GG* / 3: *CC* – 1: *GG* / *TT*

*The C allele carriers might favor power/strength-oriented sport performance* [3].

*BDNF-AS* – *rs6265*

1: *CC(Val)* / 3: *TT(Met)* – 1: *GG(Val)* / 3: *AA(Met)*

*The BDNF Met/Met genotype has a protective role in obesity in healthy subjects* [4].

*GNB3* – *rs5443*

1: *TT* / 3: *CC*

*The presence of 825T-allele may impair athletic performance and may serve as a genetic marker of low capacity for athletic performance* [5].

*DRD2 – rs1076560*

1: *TT* / 3: *GG* – 1: *AA* / 3: *CC*

*DRD2 rs1076560 T allele had a threefold increase in psychosis risk compared to GG homozygotes* [6].

*Rs1076560(A) alleles were 1.3 fold more associated with Alcoholism than the rs1076560(C allele* [7].

*SNAP-25 – rs362584*

1: *AA* / 3: *GG* (ancestral)

*SNAP-25 is synthesized in the motor nerve endings, and affects motor neurons of the spinal cord* [8].
