## Supplemental equations for "Detection of epistasis between *ACTN3* and *SNAP-25* with an insight towards gymnastic aptitude identification"

**S3 Supplementary Material:**

### Epistasis Detection in Genetic Data of Sports Gymnastics Contestants and Sedentary Individuals With an Insight Towards Classification Accuracy

This text is to give a full statistical foundation on data processing performed to verify hypotheses and answer research questions (.docx)

#### Theoretical Background – Data Analysis

**Contents:**


1. **Basic Statistics ____________________ 2**
2. **Multidimensional Statistical Description ____________________ 3**

**References ____________________ 6**

Epistasis Detection in Sportsman Recognition Context: Gymnastics Matter is intentionally expanded. The reason regards the fundamental explanation of the computational aspect of the research.

#### 1. Basic Statistics and Notations

All polymorphisms were studied through minor allele frequency and Hardy-Weinberg equilibrium according to the formula (1):

$$p^2 + 2pq + q^2 = 1, \quad (1)$$

where:

$p^2 - frequency\ of\ dominant\ homozygous\ genotype$;

$q^2 - frequency\ of\ recessive\ homozygous\ genotype$;

$2pq -\ frequency\ of\ heterogeneous\ genotype.$

In standard – linear approach genotypes were coded_as 1, 2, or 3, depending on the number of copies of a dominant allele for each SNP. Next, the most commonly used six subject-level gene models: recessive, multiplicative, additive / harmonic, dominant, and over-dominant models [1] were computed to adjust the best one to a given SNP data distribution. Each of the six models was originally established using the relationship among $odd_{MM}$, $odd_{Mm}$, and $odd_{mm}$:

the recessive model is defined by $\mathbf{odd_{Mm} = odd_{MM}}$. Therefore, OR1 = 1,

the multiplicative model is defined by $\mathbf{odd_{Mm} = \sqrt{(odd_{MM} \times odd_{mm})}}$. Hence, OR1= OR2,

the additive model is defined by $\mathbf{odd_{Mm} = (odd_{MM} + odd_{mm}) / 2}$. Hence, OR2 = 2 − 1 / OR1,

the harmonic model is defined by $\mathbf{odd_{Mm}} = 1 / ((1 / \mathbf{odd_{MM}}) + 1 / \mathbf{odd_{mm}}) / 2)$,

the dominant model is defined by $\mathbf{odd_{mm} = odd_{Mm}}$. And, OR2 = 1,

the over-dominant model is defined by $\mathbf{odd_{mm} = odd_{MM}}$. Therefore, OR2 = 1 / OR1.

After qualitative control of alleles and model selection, an information gain (IG) of every SNP was computed with standard coding and with genetic model adjustment. Briefly, IG is a reduction in entropy after partitioning sample to subgroups according to a given attribute. The larger the information gain, the more the SNP reduces uncertainty about the biological potential of sports mastery [2]. The uncertainty is consistent with Shannon entropy [3]:

$$H(S_j) = -\sum_{w=1}^{r} P_w log(P_w), \tag{2}$$

and information gain is represented by the equation:

$$IG\ (H, A) = ENTROPY\ H - \sum_{v \in A} \frac{H_v}{H} ENTROPY\ H_v. \tag{3}$$

However, statistical significance is calculated from G-square statistics. In terms of $H_0$ it expresses that the input attribute is conditionally independent with a target attribute:

$$G^2(A, D) = 2 \times \ln{(2)} \times |D| \times InfoGain(S, A), \tag{4}$$

where:

$A - target\ attribute;$

$D - alleles.$

2. **Multidimensional Data Analysis**

Considerations above comprise the first step of the framework for detecting and characterizing epistasis (gene – gene interaction). Once interesting SNPs will pass the early statistical calculation, the very next point in analysis regards the multifactor dimensionality reduction. According to Jakulin [4] concept, MDR starts from constructing interaction dendrograms under supervised clustering. For this purpose, Rajski's distance matrix was computed:

$$\langle A, B, C \rangle_R = 1 - |\bar{I}(A; B; C)|, \tag{5}$$

where:

$|\bar{I}(A; B; C)| - normalized\ interaction.$

Subsequently, Ward's method was used:

$$EES = \sum_{C_a \in N_c} \| C_a - \bar{C} \|^2, \qquad (6)$$

where:

$C_a - cluster\ a;$

$N_c - number\ of\ clusters;$

$\bar{C} - cluster\ centroid\ .$

Finally, genetic dendrogram was established by implementing Lance and Williams recursive algorithm:

$$D\big(C_i, C_j\big) = \alpha_m D(C_i, C_m) + \alpha_n D(C_i, C_n) + \beta D(C_m, C_n) + \gamma |D(C_i, C_m) - D(C_i, C_n)|, \qquad (7)$$

where:

$C_{i,j,m,n} - clusters;$

$\alpha_{m,n}, \beta, \gamma - agglomerative\ criteria;$

$D - Ward's\ distance.$

From the cognitive perspective, the very important point in epistasis overview and genetic paradigm for identification of athletes predisposed to sport is to determinate the best epistatic model. In methodology one of the most acknowledged algorithms is Relief-F:

$$W_i = W_i - \frac{\sum_{k=1}^{K} D_H}{n_c * K} + \sum_{c=1}^{C-1} P_c * \frac{\sum_{k=1}^{K} D_{M_c}}{n_c * K}, \qquad (8)$$

where:

$n_c - number\ of\ instances\ in\ class\ c;$

$D_H, D_{M_c} - difference\ function\ between\ instance R_i\ and\ H\ or\ M_c;$

$P_c - \ apriori\ probability\ for\ class\ c;$

$H - neighbors\ of\ R_i\ from\ the\ same\ class;$

$M_c - neighbors\ of\ R_i\ from\ different\ classes;$

$K - number\ of\ neighborhoods.$

The significance of result from (7) is calculated in sign test based on cross-validation consistency:

$$p = \sum_{k=c}^{n} \binom{n}{k} \left(\frac{1}{2}\right)^{n}, \qquad (9)$$

where:

$n - number\ of\ cross - validations;$

$k - number\ of\ cross - validations \geq 0{,}5.$

MDR analysis do not allow for marginal effect estimation in epistatic model. This problem is solved through multivariate logistic regression. When an outcome is categorical, the penetrance function is as follows:

$$P = p(Y = 1 \mid x_1, x_2, \dots, x_k) = \frac{\exp\,(\beta_0 + \sum_1^k \beta_i x_i)}{1 + \exp\,(\beta_0 + \sum_1^k \beta_i x_i)}, \qquad (10)$$

where:

$p(Y = 1 \mid x_1, x_2, \dots, x_k)$ – conditional probability as a function of $x_1, x_2, \dots, x_k$;

$\beta_0 - constant;$

$\beta_i - regression\ weights;$

$exp - exponentional\ function.$

The formula (9) is robust when non-linear dependencies exist between alleles. Nevertheless, linear approach for detecting an impact of genetic polymorphisms might bring several benefits:

$$\log(r/(1 - r)) = \mu + \beta_i x_i, \qquad (11)$$

where:

$r - odds\ of\ being\ successful\ athlete;$

$u - constant;$

$\beta_i - regression\ weights;$

$x_i - genotype.$

After determining marginal weights for given SNP and examining potential gene – gene interaction effect an area under the curve remains to be calculated. For this case the trapezoid was used:

$$AUC = \sum_{j=1}^{t-1}\left(x_{j+1} - x_j\right)\frac{y_{j+1}+y_j}{2}, \quad (12)$$

where:

$t - number\ of\ trapezoidal\ strips;$

$x_j - false\ positives\ classifications;$

$y_j - true\ positives\ classifications.$
