## Supplementary material for "Detection of epistasis between *ACTN3* and *SNAP-25* with an insight towards gymnastic aptitude identification": Author summary

Performance enhancing polymorphisms (PEPs) may be investigated using a univariate approach but also from an epistatic perspective. A deep understanding of the genetic foundations of athleticism is essential for scientifically-grounded procedures for diagnosis and prognosis of gymnastic aptitude. Notably, the aspect of multi-variant genetic identification of athletes is not well documented in the literature. Specifically, the cross-partial derivatives of interactions between PEPs in sports gymnastics and evaluation of pure genetic models for athlete's recognition are yet unexplored. **This study confirms the interaction between variants in the *ACTN3* and *SNAP-25* loci. The *rs1815739 x rs362584* interplay proceeds with a multiplicative – over-dominant scheme and has the potential to confer these genetic variants as diagnostic markers for athlete's discrimination. We follow up on experiments originally performed by Luigi Galvani in 1782 and 1786 but from a genetics perspective.** We found that homogenous derived genotype carriers exhibit the lowest chance of classification to the athlete group – gymnasts. Our observation of *ACTN3 * SNAP-25* interaction may be applicable in advanced talent identification procedures. The *ACTN3 – SNAP-25* interaction has never been reported before but based on the model and empirical evidence presented here, the two genes are undeniably interlinked.
