## Supplementary material for "Detection of epistasis between *ACTN3* and *SNAP-25* with an insight towards gymnastic aptitude identification": Graphical abstract

### Analysis of 7 Performance Enhancing Polymorphisms reveals epistasis between ACTN3 & SNAP-25 in the context of gymnastics

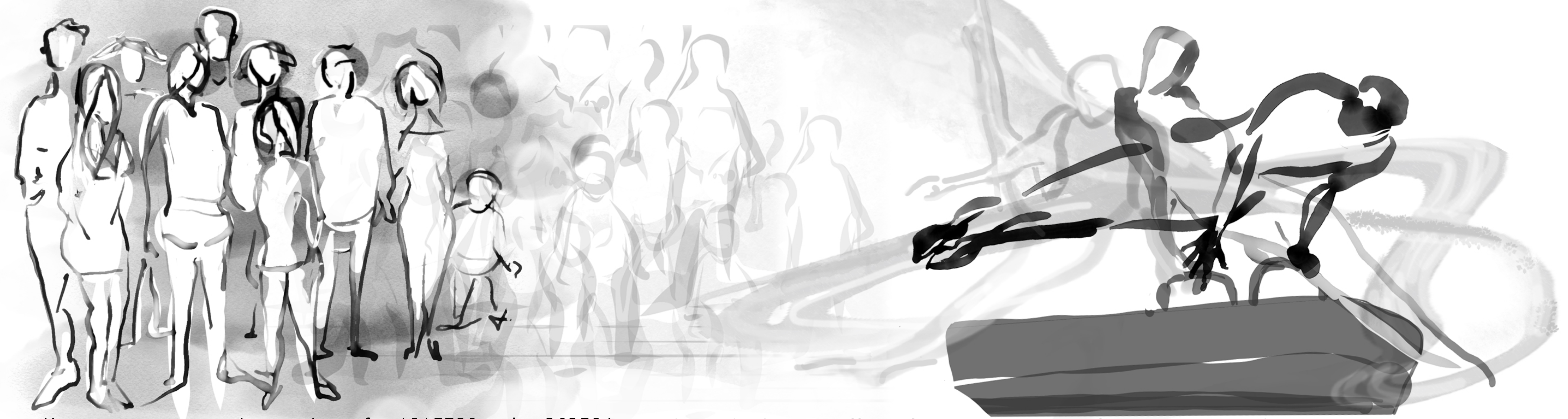

Homogeneous recessive carriers of *rs1815739* and *rs362584* have a 6% chance of becoming gymnasts

The multiplicative effect of *RR, AA* genotypes for *rs1815739* and *rs362584* increases probability of success in gymnastics to 66%.

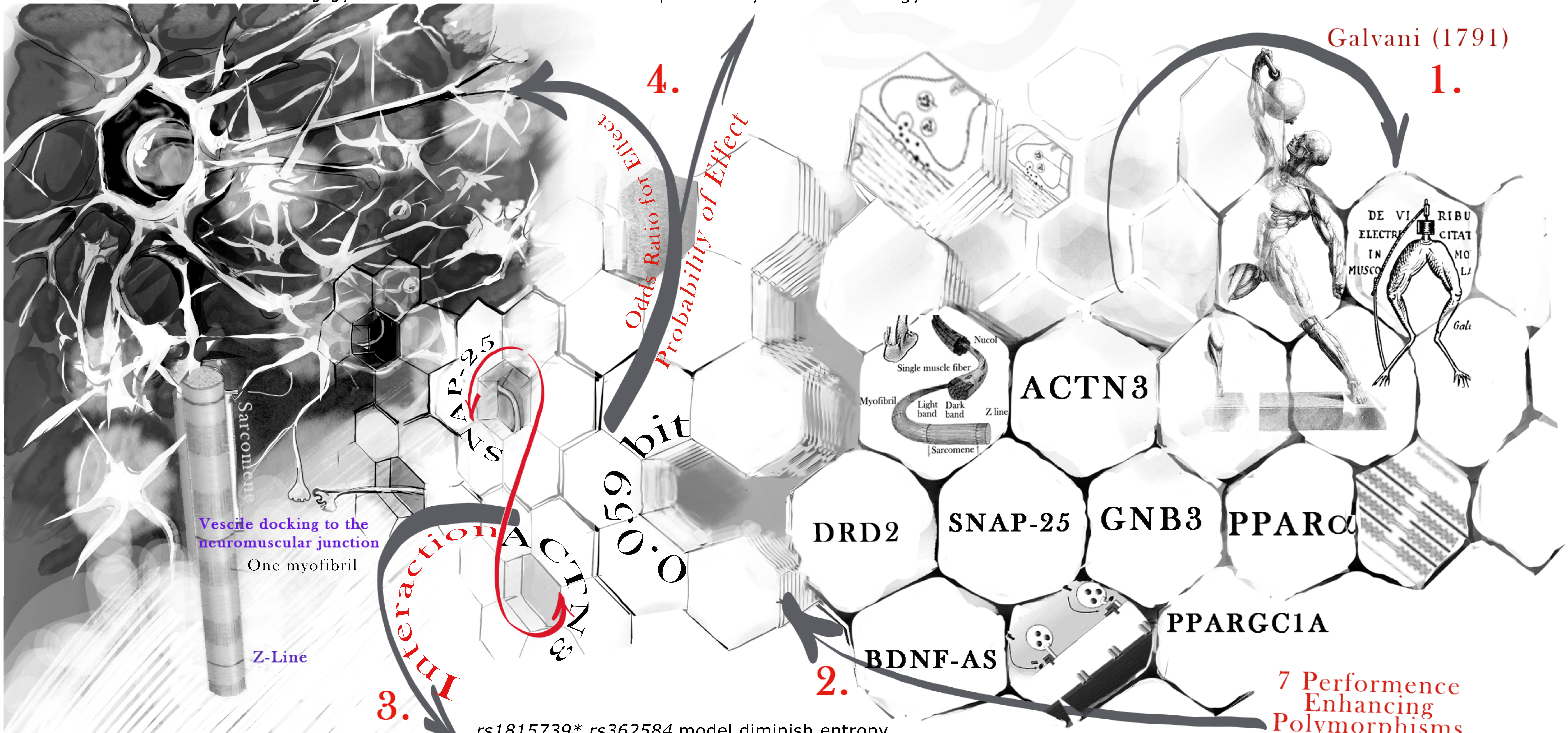


*rs1815739* rs362584* model diminish entropy of sports gymnastics (0.77 bit) in 7.66%
